# DYNC1LI2 regulates localization of the chaperone-mediated autophagy-receptor LAMP2A and improves cellular homeostasis in cystinosis

**DOI:** 10.1101/2020.06.09.142349

**Authors:** Farhana Rahman, Jennifer L. Johnson, Jinzhong Zhang, Jing He, Stephanie Cherqui, Sergio D. Catz

## Abstract

The dynein motor protein complex is required for retrograde transport but the functions of the intermediate-light chains that form the cargo-binding complex are not elucidated and the importance of individual subunits in the maintenance of cellular homeostasis is unknown. Here, using mRNA arrays and protein analysis, we show that the dynein subunit, intermediate chain 2 (DYNC1LI2) is downregulated in cystinosis, a lysosomal storage disorder caused by genetic defects in the lysosomal cystine transporter, cystinosin. Reconstitution of the expression of DYNC1LI2 in *Ctns*^-/-^ cells re-established endolysosomal dynamics. Defective vesicular trafficking in cystinotic cells was rescued by DYNC1LI2 expression which correlated with decreased endoplasmic reticulum stress manifested as decreased expression levels of the chaperone Grp78. Mitochondrial fragmentation in cystinotic fibroblasts was also rescued by DYNC1LI2. Survival of cystinotic cells to oxidative stress insult was increased by DYNC1LI2 reconstitution but not by its paralog DYNC1LI1, which also failed to decrease ER stress levels and mitochondrial fragmentation. Restoring DYNC1LI2 expression rescued the localization of the chaperone-mediated autophagy receptor, LAMP2A, and restored cellular homeostasis of cystinotic proximal tubule cells, the primary cell type affected in cystinosis. DYNC1LI2 failed to rescue phenotypes in cystinotic cells when LAMP2A was downregulated or when co-expressed with dominant negative (DN) RAB7 or DN-RAB11, which impair LAMP2A trafficking. DYNC1LI2 emerges as a new target to repair underlying trafficking and CMA defects in cystinosis, a mechanism that is not restored by currently used lysosomal depletion therapies.

## INTRODUCTION

Cystinosis is a lysosomal storage disorder caused by the accumulation of the amino acid cystine due to genetic defects in the *CTNS* gene, which encodes cystinosin, the lysosomal cystine transporter^1,2^. Increased levels of intra-lysosomal cystine lead to cell malfunction, which is specially manifested in the kidney’s epithelial proximal tubule cells (PTCs). However, many cellular defects in cystinosis are independent of lysosomal overload^3–5^, suggesting that cystinosin regulates cellular processes independently of its role as a cystine transporter. The efficiency of cysteamine in retarding the rate of glomerular deterioration and improvement of linear growth in children with cystinosis demonstrates the effectiveness of cystine-depleting therapies^6–8^. However, even with cysteamine treatment, cell malfunction, tissue failure and progressive renal injury still occurs^9^, thus, cystine accumulation is not the only cause of all the defects observed in cystinosis^9,10^. Although the molecular defects that initiate or promote disease progression in cystinosis are not fully characterized, several evidences point to defects in vesicular trafficking as one of the molecular mechanisms that are defective in cystinosis but not repaired by cysteamine^11,12^.

Dynein, a large motor protein complex, carries out an extensive range of cellular processes. The dynein family includes axonemal dyneins that drive ciliary and flagellar motility^13^ and cytoplasmic dyneins which perform several intracellular processes including cargo transport, organelle trafficking and mitotic spindle assembly and positioning^14,15^. Dynein controls cellular functions through regulatory movement to the minus-end of the microtubules, a process referred to as retrograde transport. Cytoplasmic dynein complexes are composed of dynein intermediate chains, light-intermediate chains (LIC) and light chains, which form a complex with two identical subunits of dynein heavy chains^16^. The complex is regulated by dynactin and adaptor proteins^17–19^, which facilitate the movement along microtubules^20^, but how dynein light chain subunits help perform individual cellular functions is not completely understood.

Individual dynein subunits may allow specific functions and interactions with specific binding partners that regulate dynein motor activity^18^. While the dynein heavy chain polypeptides provide the energy that facilitates movement along microtubules, the functional roles of the intermediate, light-intermediate (LIC) and light chain polypeptides remain poorly understood. The dynein LIC subunits present in cytoplasmic dyneins are proposed to regulate cargo attachment and dynein transport^21–24^. The two genes which encode LICs (DYNC1LI1 and DYNC1LI2) showed 63% sequence identity in humans and share high sequence similarity within the dynein- and cargo-binding domains^25,26^.

LIC1 and LIC2 were found to perform overlapping functions in cells in earlier studies^27^. However, a recent work demonstrated that both LICs have distinct roles in neurogenesis, migration and translocation^25,28^. Mammalians LICs bind to GDP *via* their Ras-like G-domain, which may be released upon binding to the dynein heavy chain^29^. Furthermore, the C-terminal domain of LIC1 has been shown to bind directly to RILP^30^, Rab11-FIP3^31,32^ and BicD2^18,33^, which are effector molecules of the Rab GTPases Rab7, Rab11 and Rab6, respectively. Thus, LICs bridge the motor heavy chain to specific vesicle-associated molecular effectors, and thus may contribute to the control of the specificity of cargoes during retrograde transport.

Autophagy is a regulated cellular mechanism of degradation of cytoplasmic material in lysosomes, that contributes to nutrient preservation and maintains cellular homeostasis. Chaperone-mediated autophagy (CMA)^34^ is a selective form of autophagy consisting of the chaperone-dependent selection of soluble cytosolic proteins with a KFERQ-like signal peptide that are then targeted to lysosomes and directly translocated across the lysosome membrane for degradation^35^. The targeted substrate-chaperon complex binds to the lysosomal surface through the cytosolic tail of the lysosomal-associated membrane protein type 2A (LAMP2A)^36^, the only CMA receptor. Similar to macroautophagy, CMA upregulation occurs during prolonged nutritional stress (starvation), exposure to toxic compounds, and mild oxidative stress^37^ suggesting a role for this pathway in the selective removal of abnormal or damaged proteins under these conditions^38^. In previous studies, we revealed a defective mechanism of CMA in cystinosis^3,39,40^. In particular, we showed defective trafficking and mislocalization of the CMA receptor LAMP2A in cystinosis, a process associated with impaired cellular homeostasis manifested as increased endoplasmic reticulum stress and decreased survival to oxidative stress^3^. Furthermore, we established that correct LAMP2A localization depends on trafficking mechanisms regulated by the small GTPases Rab11 and Rab7^40^.

Here, to better understand the mechanisms underlying vesicular transport defects in cystinosis, we used an mRNA array approach and identified the motor protein subunit dynein light intermediate chain DYNC1LIC2 to be downregulated in cystinosis. Rescue of DYNC1LI2 expression increases LAMP2A trafficking and improves cell homeostasis in both cystinotic fibroblast and proximal tubule epithelial cells. Mechanistically, the rescue of cystinotic cell function by DYNC1LI2 requires functional Rab11 and Rab7 GTPases, and the process is inhibited by downregulation of LAMP2A. Thus, DYNC1LIC2 regulates vesicular trafficking and lysosomal-associated protein sorting through a mechanism control by chaperone-mediated autophagy and is essential to restore cellular homeostasis in cystinosis.

## RESULTS

### DYNC1LI2 is downregulated in cystinotic cells

Vesicular trafficking is defective in cystinosis^12^. To identify potential molecular regulators of vesicular trafficking that are dysregulated in cystinosis, we performed mRNA analysis of cystinotic kidneys. DYNC1LIC2, a light intermediate subunit of the motor protein dynein was identified to be downregulated in six independent cystinosis kidney samples (p<3.32×10^−5^). DYNC1LI2 downregulation at the protein level in cystinosis was confirmed by immunoblotting of WT and *Ctns*^-/-^ mouse embryonic fibroblasts (MEFs) (**Fig. 1A**). Quantitative immunoblotting analysis shows that DYNC1LI2 expression is significantly decreased in cystinotic cells compared to wildtype cells (**Fig. 1B**). As a control, we also examined the expression of the DYNC1LI1 subunit, a paralog of DYNC1LI2. Different from DYNC1LI2, the expression of DYNC1LI1 in *Ctns*^-/-^ cells was not different from that observed in wild-type cells (**Figs. 1C and D**). The selective decreased expression of DYNC1LI2 was also confirmed using an independent approach consisting of the immunofluorescence analysis of endogenous DYNC1LI2 protein (**Figs. 1E and F**). Finally, we engineered CRISPR-Cas9 cystinotic human proximal tubule cells (*CTNS*-KO PTCs)^39^ which, similar to *Ctns*^-/-^ fibroblasts, express significantly decreased levels of DYNC1LI2 (**Figs. 1G and H**), but normal levels of the paralog DYNC1LI1 (**Figs. 1G and I**).

**Figure 1.**
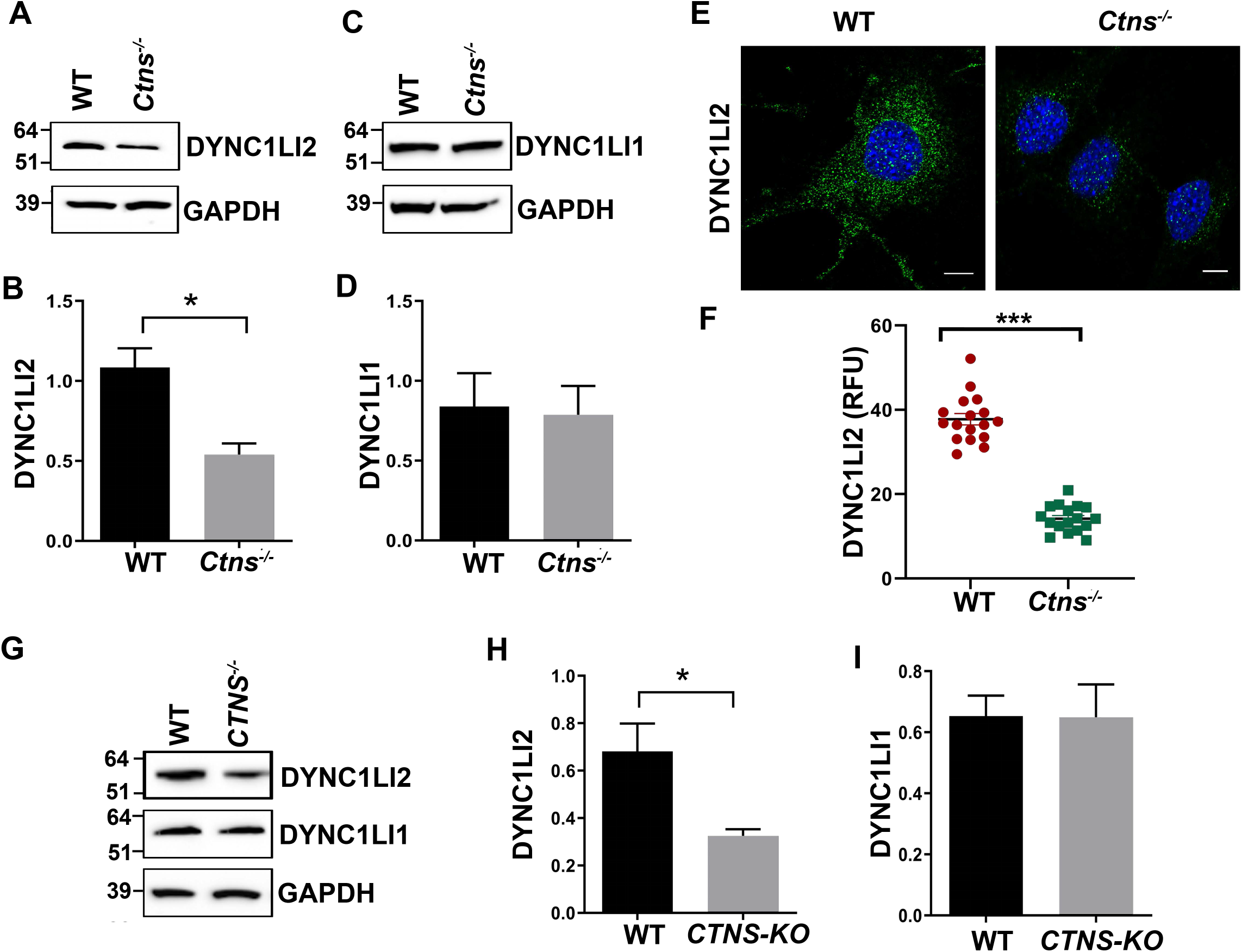
DYNC1LI2 but not DYNC1LI1 is downregulated in cystinotic fibroblasts and CTNS-KO proximal tubule cells (PTCs) *A*, Expression of DYNC1LI2 in WT and *Ctns*^-/-^ fibroblasts was analyzed by Western blot. *B*, Quantification of DYNC1LI2 expression levels from 3 independent experiments. *C*, Expression of DYNC1LI1 in WT and *Ctns*^-/-^ mouse fibroblasts analyzed by Western blot. *D*, Quantification of DYNC1LI1 expression levels from 3 independent experiments. *E*, Immunofluorescence analysis of DYNC1LI2 expression in wild-type and *Ctns*^-/-^ MEFs. *Scale bar*, 10 μm. *F*, Quantification of DYNC1LI2 fluorescence intensity (RFU). The data correspond to 3 independent experiments. *G*, Expression of DYNC1LI2 and DYNC1LI1 in WT and *CTNS-KO* human PTCs analyzed by Western blot. *H and I*, Quantification of DYNC1LI1 and DYNC1LI2 expression levels from 3 independent experiments. Mean ± SEM *, *p*< 0.05, ***, *p*< 0.001.

### DYNC1LI2 rescues vesicular trafficking and the localization of the CMA receptor, LAMP2A, in cystinosis

Cystinotic cells are characterized by impaired vesicular trafficking mechanisms that have deleterious consequences on cell homeostasis and function^12,40^. In this work, to analyzed whether reconstitution of DYNC1LI2 expression rescues the vesicular transport impaired phenotypes and associated defects in cystinotic cells, we stably reconstituted DYNC1LI2 expression or, as control, stably expressed DYNC1LI1 in *Ctns*^-/-^ cells by lentiviral infection. Next, to analyze the impact of DYNC1LI2 reconstitution on vesicular dynamics in *Ctns*^-/-^ cells, we first studied endolysosomal trafficking in cells labeled with the acidotropic dye LysoTracker, by Total Internal Reflection Fluorescence microscopy (TIRFM). We found that the dynamics of vesicles in *Ctns*^-/-^ cells was significantly impaired compared to wild-type cells (**Figs. 2A and B**). In these assays, the vesicles were segregated according to their speed, and *Ctns*^-/-^ cells showed increased numbers of non-motile vesicles and vesicles moving at very low speed with a concomitant decreased of fast-moving vesicles (**Fig. 2B**). The defective phenotype was rescued by reconstitution of DYNC1LI2-expression. Next, we asked whether the exogenous expression of the paralog DYNC1LI1 also improves endolysosomal trafficking in these cells. Exogenous expression of DYNC1LI1 also rescued the defective trafficking phenotype in *Ctns*^-/-^ cells albeit not as efficiently as DYNC1LI2. Altogether the data support the premise that expression of DYNC1LI1/2 in *Ctns*^-/-^ fibroblasts re-establishes endolysosomal dynamics.

**Figure 2.**
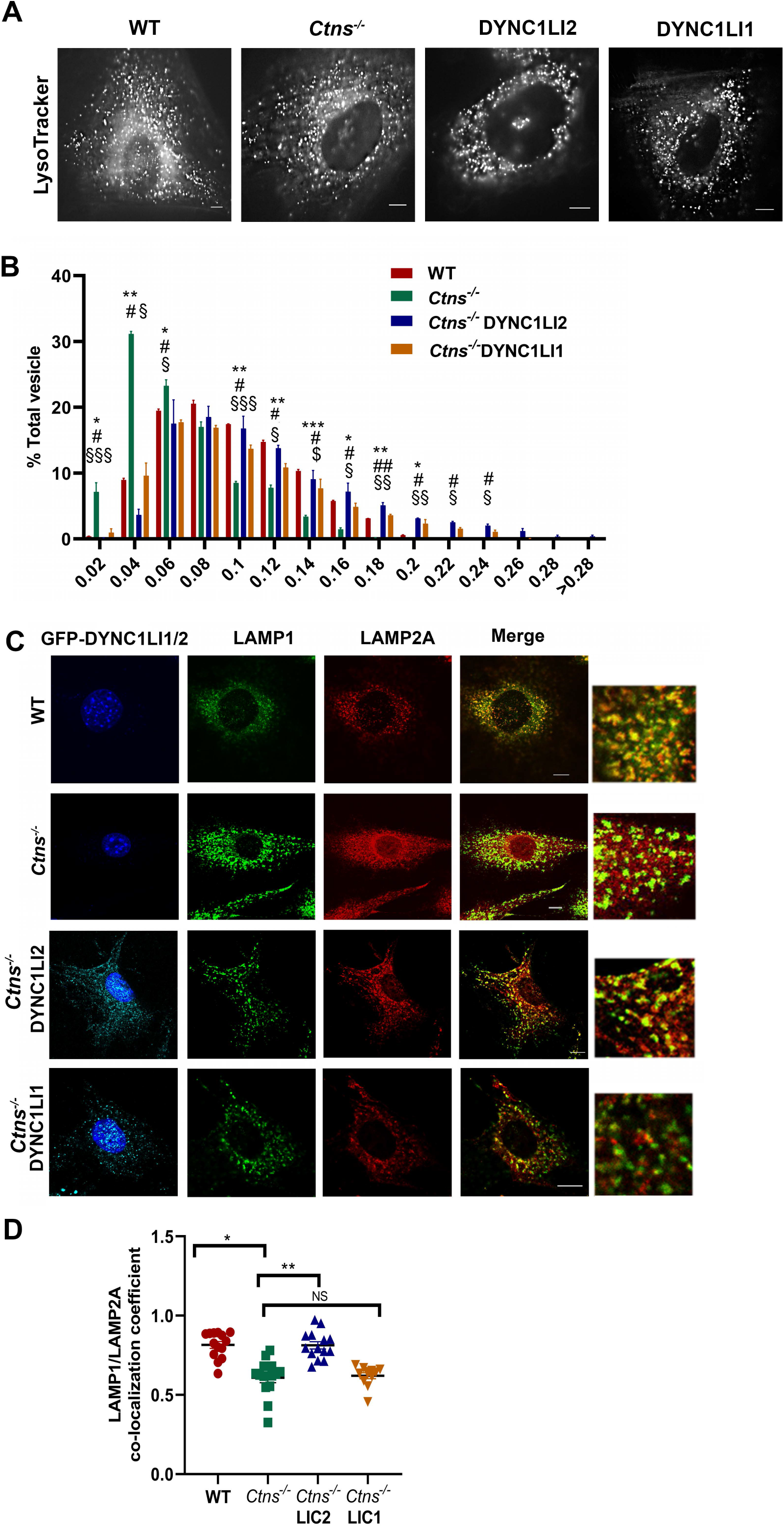
DYNC1LI2 rescues vesicular trafficking and lysosomal localization of LAMP2A in cystinotic cells. *A*, Representative images from the analysis of vesicle movement in wild-type, *Ctns*^-/-^, DYNC1LI2- and DYNC1LI1-expressing *Ctns*^-/-^ cells by TIRFM. Scale bar: 10 μm. *B*, the speeds for the independent vesicles were binned in 0.02 μm/s increments and plotted as a percentage of total vesicles for a given cell. The statistically significant differences between the groups are indicated in the figure. Mean ± SEM *, *p*< 0.05, **, *p*< 0.01, ***, *p*< 0.001; WT *vs Ctns*^-/-^, ^#^, *p*< 0.05, ^# #^, *p*< 0.01, *Ctns^-/-^ vs* DYNC1LI2 and §, *p*< 0.05, §§, *p*< 0.01, §§§, *p*< 0.001; *Ctns^-/-^ vs* DYNC1LI1. *C*, Immunofluorescence analysis of endogenous LAMP1 (pseudo-colored green) and LAMP2A (red) localization in WT, *Ctns*^-/-^, GFP-(pseudo-colored cyan)-DYNC1LI2-, and GFP-DYNC1LI1-expressing *Ctns*^-/-^ cells. *Scale bar*, 10 μm. *D*, LAMP2A/LAMP1 colocalization increases in response to GFP-DYNC1LI2 expression. Mean ± SEM, *, *p*< 0.05; **, *p*< 0.01; NS, not significant.

Defective CMA in cystinosis is associated with impaired LAMP2A trafficking and the CMA receptor is retained in Rab11-positive carrier vesicles in cystinotic cells, a phenotype that is rescued by the expression of constitutively active Rab11 but not by cysteamine treatment^3,39,40^. Here, we next analyzed whether DYNC1LI2 or DYNC1LI1 were able to rescue the mislocalization of the CMA receptor, LAMP2A, at lysosomes. To this end, we performed confocal microscopy analysis of the localization of LAMP2A in relation to the lysosomal marker LAMP1. We show a significantly increased co-localization of the CMA receptor LAMP2A with LAMP1-positive puncta in *Ctns*^-/-^ cells after reconstitution of DYNC1LI2 expression. Contrarily, DYNC1LI1 over-expression did not rescue the LAMP2A-mislocalized phenotype (**Figs. 2C and D**), despite its partial rescue effect on endolysosomal trafficking (**Fig. 2A and B**). Altogether, these results suggest DYNC1LI2 but not DYNC1LI1 regulates the trafficking of LAMP2A-carrying intermediate vesicles^3^, thus establishing unique mechanistic and trafficking properties for DYNC1LI2 in this cellular system.

### DYNC1LI2 but not DYNC1LI1 decreases ER stress

In cystinosis, CMA defects are associated with increased endoplasmic reticulum (ER) stress and susceptibility to oxidative stress^3,12^. To evaluate whether the rescue of DYNC1LI2 expression has a positive impact on cellular homeostasis in cystinosis, we next analyzed the effect of DYNC1LI2 reconstitution on the expression levels of the UPR target gene Grp78, in *Ctns*^-/-^ fibroblasts. Immunoblotting analysis showed that DYNC1LI2 reconstitution induces a significant decreased in the expression of endogenous Grp78 in *Ctns*^-/-^ fibroblasts compared to control cystinotic cells (**Figs. 3A and B**). Different from DYNC1LI2, exogenous expression of DYNC1LI1 failed to decrease ER stress levels in *Ctns*^-/-^ fibroblast (**Figs. 3C and 3D**).

**Figure 3.**
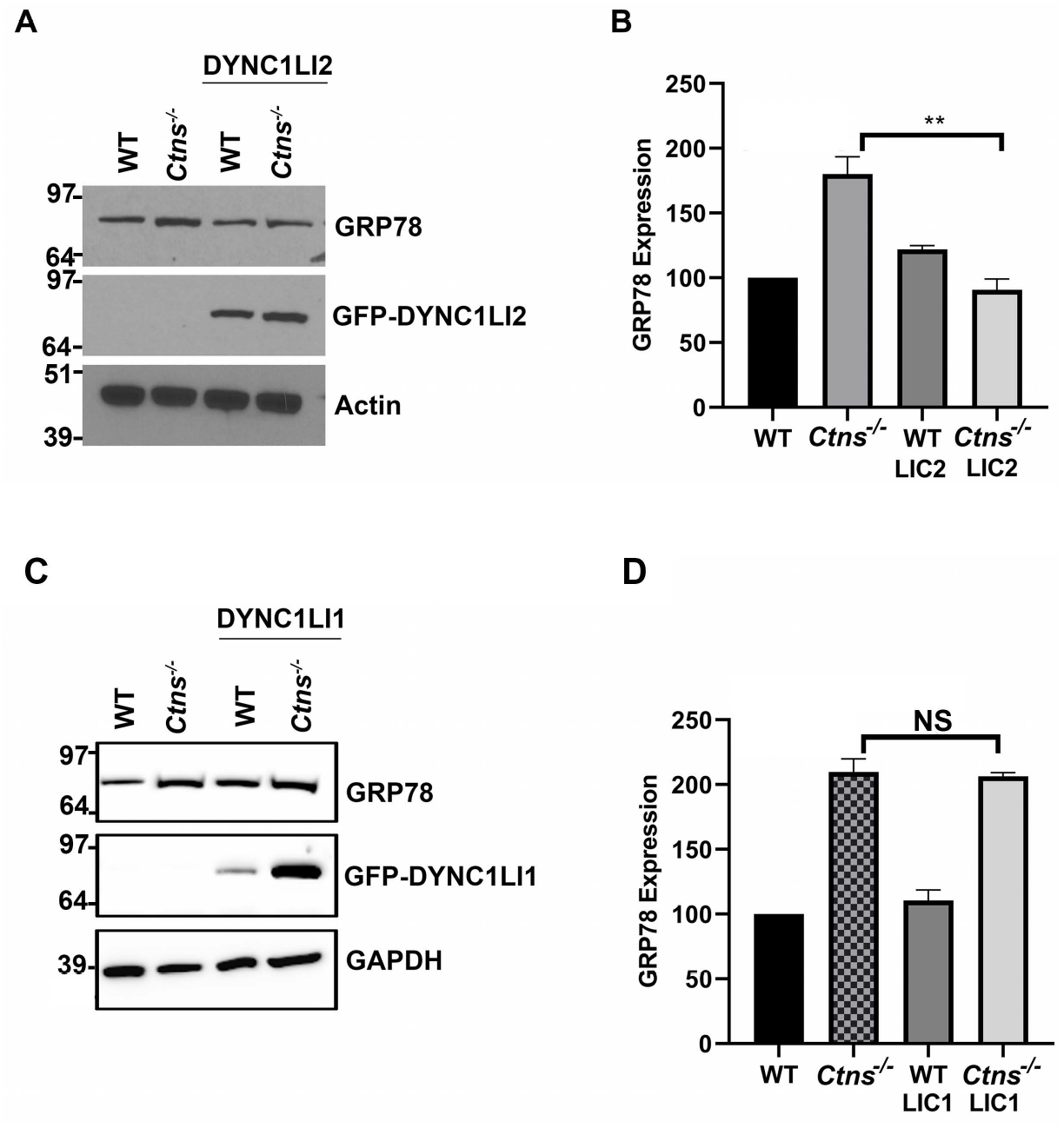
Rescue of DYNC1LI2 expression reduces ER stress in Ctns^-/-^ cells. *A*, The expression level of the established UPR target gene GRP78, which is induced during conditions of ER stress, was analyzed in wild-type (WT), *Ctns*^-/-^, and GFP-DYNC1LI2-expressing WT and *Ctns*^-/-^ fibroblasts by Western blotting, using a mouse monoclonal anti-KDEL antibody. *B*, Quantitation of GRP78 expression levels from 3 independent experiments. *C*, GRP78 expression in wild-type, *Ctns*^-/-^, and GFP-DYNC1LI1-expressing WT and *Ctns*^-/-^ fibroblasts by Western blot. D, Quantification of GRP78 expression levels from 3 independent experiments. Mean ± SEM, **, *p*< 0.01; NS, not significant.

### DYNC1LI2 decreases mitochondria fragmentation and protects cystinotic cells from oxidative stress-induced cell death

Mitochondrial fission is an important mechanism that precedes elimination of damaged mitochondrial by mitophagy^41^ and mitochondrial fragmentation is induced by oxidative stress and has been associated with human disease. Mitochondrial dysfunction has long been demonstrated in many LSDs including cystinosis^42^. Here, to establish the impact of DYNC1LI2 reconstitution on mitochondrial fragmentation in cystinotic cells, we measured mitochondria fragmentation in wild-type and *Ctns*^-/-^ cells expressing DYNC1LI2 or DYNC1LI1. We found that the average length of mitochondria is significantly longer in wild-type cells, reaching lengths double that of the average mitochondrial length in *Ctns*^-/-^ cells (**Figs. 4 A and B**). Remarkably, mitochondrial length in *Ctns*^-/-^ cells was significantly increased by the expression of DYNC1LI2 but not DYNC1LI1 (**Fig. 4B**) indicating that DYNC1LI2-specific trafficking mechanisms rescue the mitochondrial morphology-defective phenotype in cystinosis.

**Figure 4.**
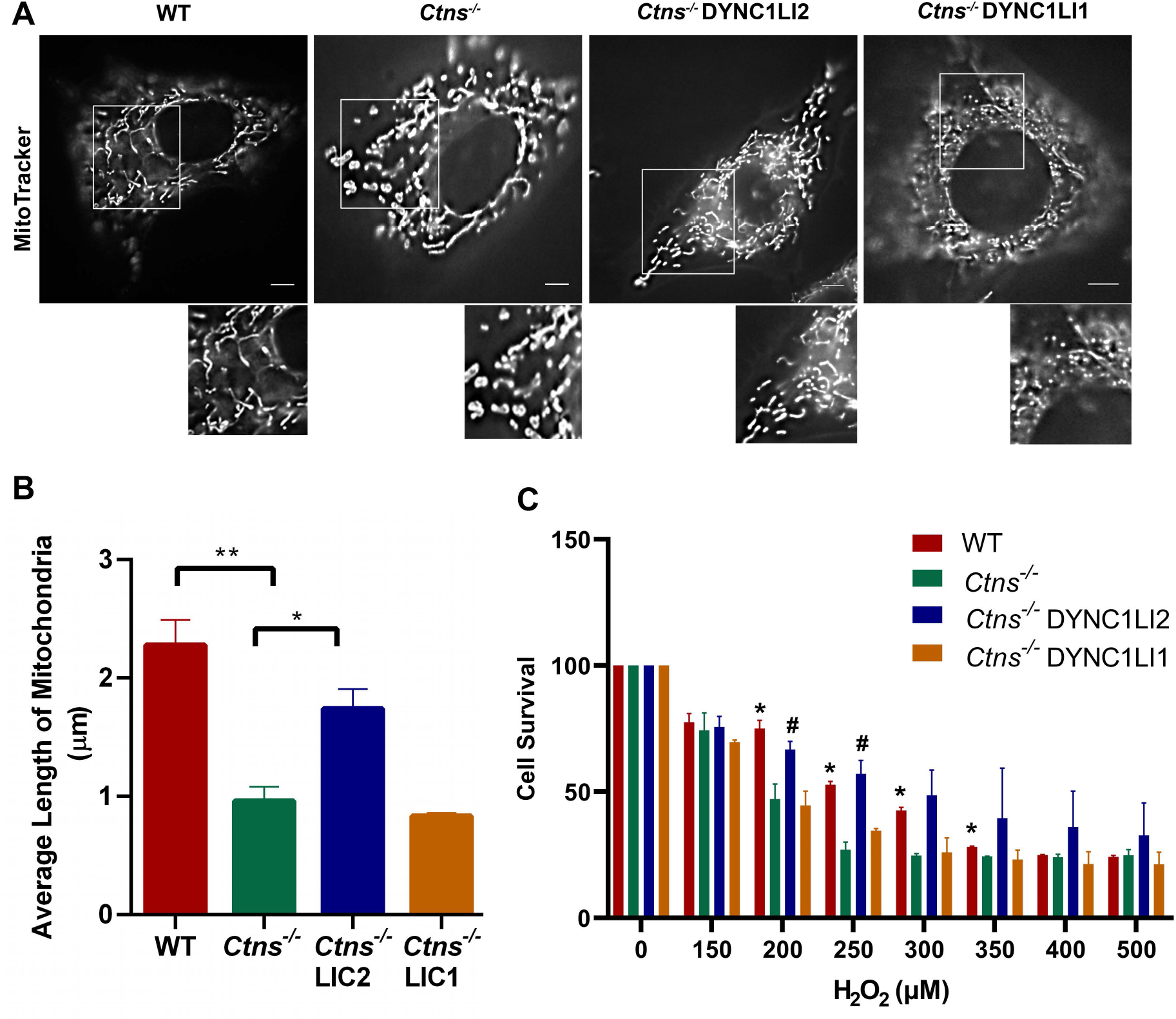
Reconstitution of DYNC1LI2 but not DYNC1LI1 expression decreases mitochondrial fragmentation and increases cell survival of Ctns^-/-^ cells. *A*, Analysis of mitochondrial length in wild-type (WT), *Ctns*^-/-^, DYNC1LI2- and DYNC1LI1-expressing *Ctns*^-/-^ cells by pseudo-TIRFM (oblique illumination) using MitoTracker. *Scale bar*, 6 μm. *B*, Quantification of mitochondrial length. Mean ± SEM, *, *p*< 0.05; **, *p*< 0.01 n=3, 8-10 cells per experiment. *C*, GFP-DYNC1LI1 and GFP-DYNC1LI2-reconstituted MEFs were treated with H_2_O_2_ and cell viability was analyzed by the MTT assay. Mean ± SEM, *, *p*< 0.05, WT *vs Ctns*^-/-^; ^#^, *p*< 0.05, *Ctns^-/-^ vs* DYNC1LI2.

Because mitochondrial dynamics have direct impact on cell death^43^ and impaired CMA leads to increased susceptibility to oxidative stress-induced cell death in cystinotic cells^3^ while CMA upregulation increases survival, we next analyzed the effect of DYNC1LI2 on cell survival to prooxidant insults. To this end, wild-type, and *Ctns*^-/-^ cells expressing either DYNC1LI2 or DYNC1LI1 were exposed to increasing concentrations of H2O2 and the cellular metabolic activity was analyzed by the MTT assay as an indication of cell survival. *Ctns*^-/-^ cells showed increased susceptibility to oxidative stress-induced cell death compared to wild-type cells. Notably, the expression of DYNC1LI2 was protective, significantly increasing the survival of *Ctns*^-/-^ cells (**Fig. 4C**) and this positive effect was not observed in cells expressing the paralog light intermediate subunit, DYNC1LI1.

### DYNC1LI2 corrects the cellular homeostasis-defective phenotypes of cystinotic proximal tubule cells

Proximal tubular cells (PTCs) are the most affected cells in nephropathic cystinosis and the loss of their apical receptors, including megalin, leads to PTC dedifferentiation and contribute to the development of Fanconi syndrome^44^. Similar to fibroblasts, cystinotic PTCs have decreased DYNC1LI2 expression (**Fig. 1**), LAMP2A mislocalization, defective CMA and impaired vesicular trafficking^11,39^. Furthermore, *CTNS-KO* cells are characterized by increased GRP78 expression and ER stress and decreased expression of the Rab-GTPase Rab11, a carrier of the chaperone-mediated autophagy receptor LAMP2A and a regulator of megalin trafficking to the apical membrane^11,39^. To study the possible beneficial impact of DYNC1LI2 expression reconstitution in PTC function, we next analyzed the localization of the CMA receptor LAMP2A in DYNC1LI2-expressing cystinotic PTCs (*CTNS*-KO) which were developed using CRISPR-*Cas*9 technology^39^. We found that DYNC1LI2 reconstitution rescues LAMP2A localization at lysosomes in *CTNS*-KO cells (**Figs. 5A and B**). Furthermore, DYNC1LI2 corrected vesicular trafficking (**Figs. 5C and D**), in particular, DYNC1LI2 recovered the dynamic behavior of a subpopulation of highly motile vesicles, those moving at >0.2μm/sec, which were previously associated with Rab7-dependent movement^11^. In addition, DYNC1LI2 decreased ER stress levels (**Figs. 5E and F**) and partially corrected the mitochondria hyper-fragmentation phenotype in cystinotic PTCs (**Figs. 5G and H**).

**Figure 5.**
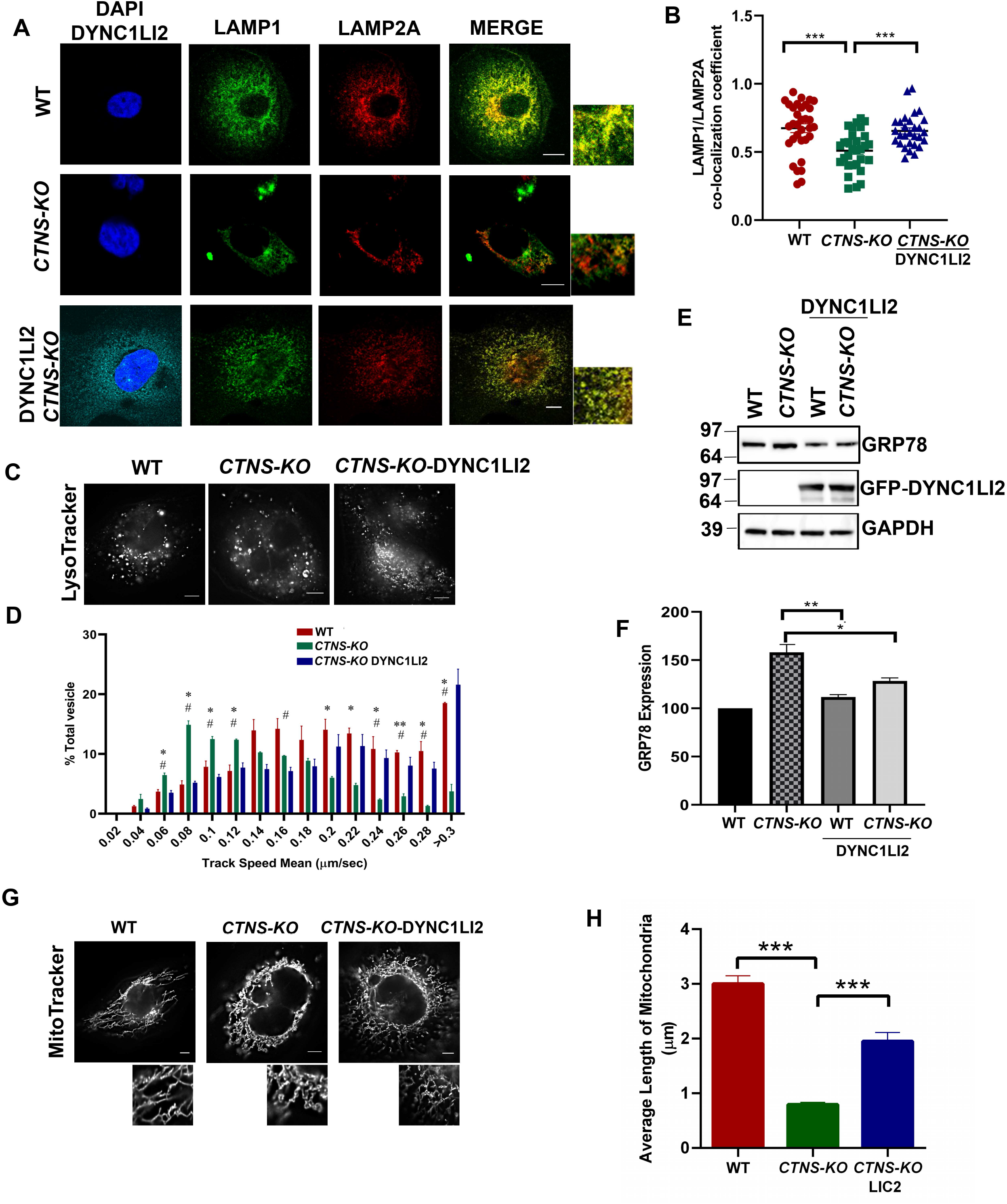
DYNC1LI2 rescues cellular homeostasis phenotypes in cystinotic proximal tubule cells (PTCs) *A*, Immunofluorescence analysis of endogenous LAMP1 (pseudo-colored green) and LAMP2A (red) localization in wild-type, *CTNS-KO* and GFP (pseudo-colored cyan)-DYNC1LI2-expressing *CTNS*^-/-^ PTCs cells. *Scale bar*, 10 μm. *B*, Quantification of LAMP2A/LAMP1 colocalization from three independent experiments. *C*, Pseudo-TIRF microscopy analysis of PTCs labeled with LysoTracker. *Scale bar*, 6 μm *D*, Analysis of vesicle movement of wild-type, *CTNS-KO* and DYNC1LI2 expressing *CTNS-KO* cells. The speeds for the independent vesicles were binned in 0.02 μm/s increments and plotted as a percentage of total vesicles for a given cell. Results are represented as mean ± SEM from at least 10 cells per experiments form 3 independent experiments. The statistically significant differences between the groups are indicated in the figure. Mean ± SEM, *, *p*< 0.05, **, *p*< 0.01, WT *vs CTNS-KO*^-^ and **^#^**, *p*< 0.05, *CTNS-KO vs CTNS-KO* DYNC1LI2. *E*, GRP78 expression in wild-type, *CTNS-KO*, and GFP-DYNC1LI2-expressing WT and *CTNS-KO* PTCs by Western blotting. *F*, Quantification of GRP78 expression from 3 independent experiments. *G*, TIRF microscopy analysis of PTC labeled with MitoTracker. *Scale bar*, 6 μm. *H*, Quantification of mitochondrial length of DYNC1LI2-expressing *CTNS-KO* PTCs n=2, 10-12 cells. B, F and H, Mean ± SEM *, *p*< 0.05, **, *p*< 0.01, ***, *p*< 0.001.

### DYNC1LI2-mediated rescue of cystinotic cell function requires both Rab11 and Rab7 but not Rab5

DYNC1LI1 and DYNC1LI2 function as linkers between the motor subunit and the transported organelle through interactions with multiple regulatory molecules^18,33^. Two of these molecules, RILP and Rab11-FIP3, are specific effectors of the small Rab GTPases Rab7 and Rab11, which are also known to regulate late and recycling endosomal trafficking, respectively. Coincidently these GTPases mediate LAMP2A trafficking and are therefore important regulators of CMA^11^. Although both DYNC1LI1 and DYNC1LI2 are described to bind to RILP and Rab11-FIP3, differential regulation of these interactions by PKA-mediated phosphorylation led to the suggestion that DYNC1LI2 regulates a subpopulation of dynein-associated endolysosomes different from that regulated by DYNC1LI1^45^.

To analyze whether the rescue mediated by DYNC1LI2 requires Rab GTPases of the endocytic pathways, we analyzed homeostatic mechanisms in cystinotic cells reconstituted with DYNC1LI2 and expressing dominant-negative (DN) Rab5, Rab7 or Rab11 GTPases, mutants that are only present in the GDP-bound form and are therefore unable to bind to their specific effectors. The cells were utilized in ER stress analysis by two independent approaches. First, we performed immunofluorescence analysis of endogenous KDEL-positive ER chaperones (**Figs. 6A and B**). Second, we analyzed the expression level of GRP78 by immunoblot (**Figs. 6C and D**). In both assays, we show that DN-Rab7 and DN-Rab11 but not DN-Rab5 preclude the rescue of the phenotype mediated by DYNC1LI2. Thus, using two independent assays, we found that the rescue of the increased ER stress phenotype mediated by DYNC1LI2 requires both functional Rab7 and Rab11 but not Rab5 GTPases.

**Figure 6.**
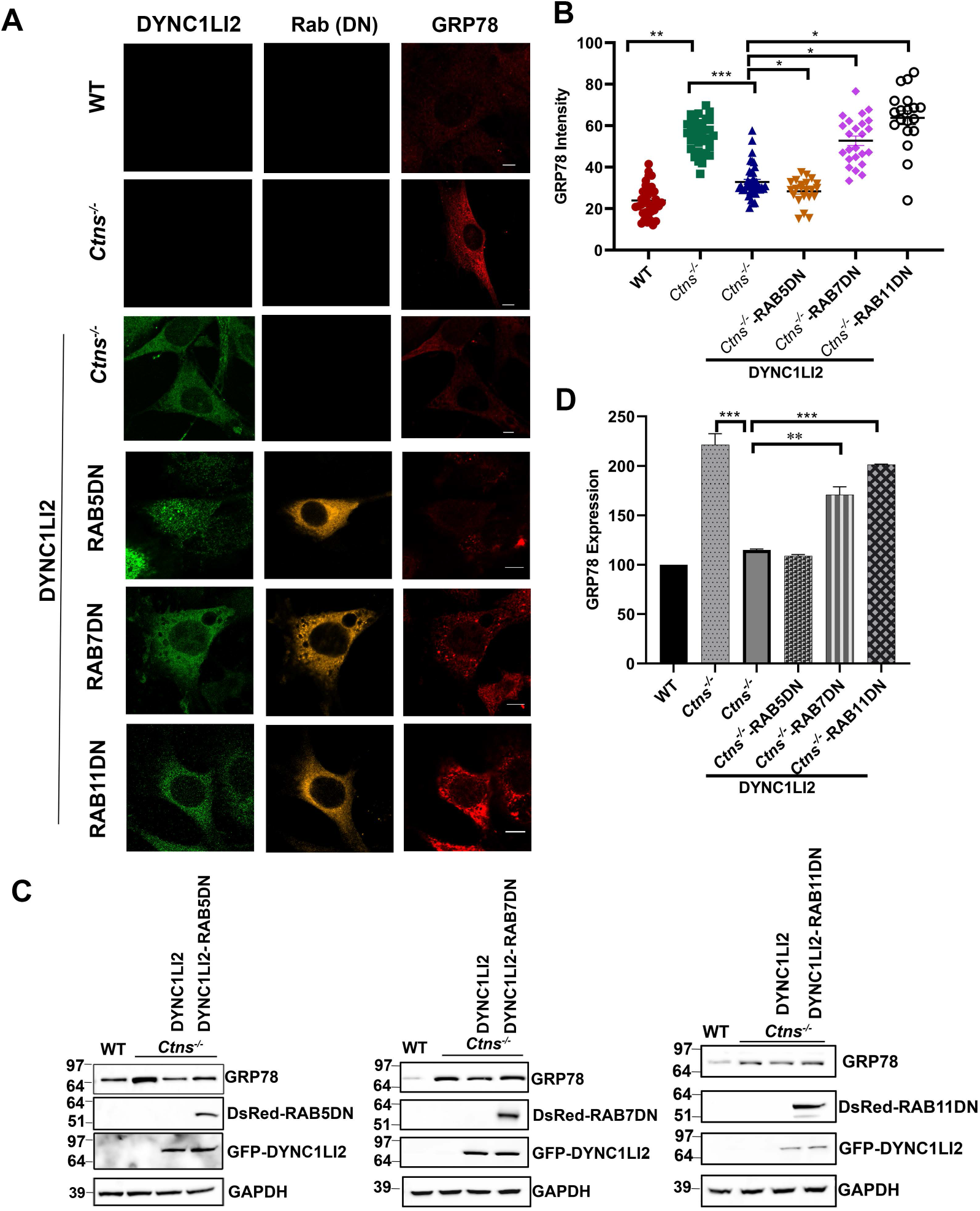
DYNC1LI2-dependent rescue of ER stress requires functional RAB7 and RAB11 but not RAB5. A, Immunofluorescence analysis of the endogenous ER stress marker, GRP78 (Red), expression in wild-type, *Ctns*^-/-^, *Ctns*^-/-^-DYNC1LI2 and dominant negative (DN)-expressing RAB5DN, RAB7DN, RAB11DN, *Ctns*^-/-^-DYNC1LI2 cells. GFP-DYNC1LI2 is shown in green. DN-Rab GTPases are pseudocolored orange. *Scale bar*, 10 μm. *B*, Quantification of ER stress marker expression from three independent experiments. Mean ± SEM, *, *p*< 0.05, **, *p*< 0.01, ***, *p*< 0.001. C, *Analysis of* GRP78 expression by Western blot in WT, *Ctns*^-/-^, *Ctns*^-/-^-DYNC1LI2 or cells expressing DN-Rabs. *D*, Quantitation of ER stress expression levels by immunoblot from 3 independent experiments. Mean ± SEM, **, *p* < 0.01, ***, *p* < 0.001.

Next, we studied whether functional Rab GTPases were necessary for DYNC1LI2 to rescue mitochondrial fragmentation and vesicular trafficking. To this end, we first analyzed mitochondrial morphology in Rab5DN, Rab7DN or RAB11DN-expressing *Ctns*^-/-^ cells reconstituted with DYNC1LI2. We found that DYNC1LI2 was able to rescue mitochondrial dynamics when coexpressed with Rab5DN but not when Rab7DN or Rab11DN were co-expressed in these cells (**Figures 7A and B**). Also, the upregulation of DYNC1LI2 in *Ctns*^-/-^ cells failed to re-establish normal endolysosomal dynamics when co-expressed with non-functional endolysosomal Rabs, Rab7DN and RAB11DN, (**Figures 7C and D**). Rab5DN only weakly interfered with DYNC1LI2-mediated trafficking upregulation, thus establishing Rab7 and Rab11 as the main effectors of DYNC1LI2-mediated upregulation of vesicular trafficking in cystinosis.

**Figure 7.**
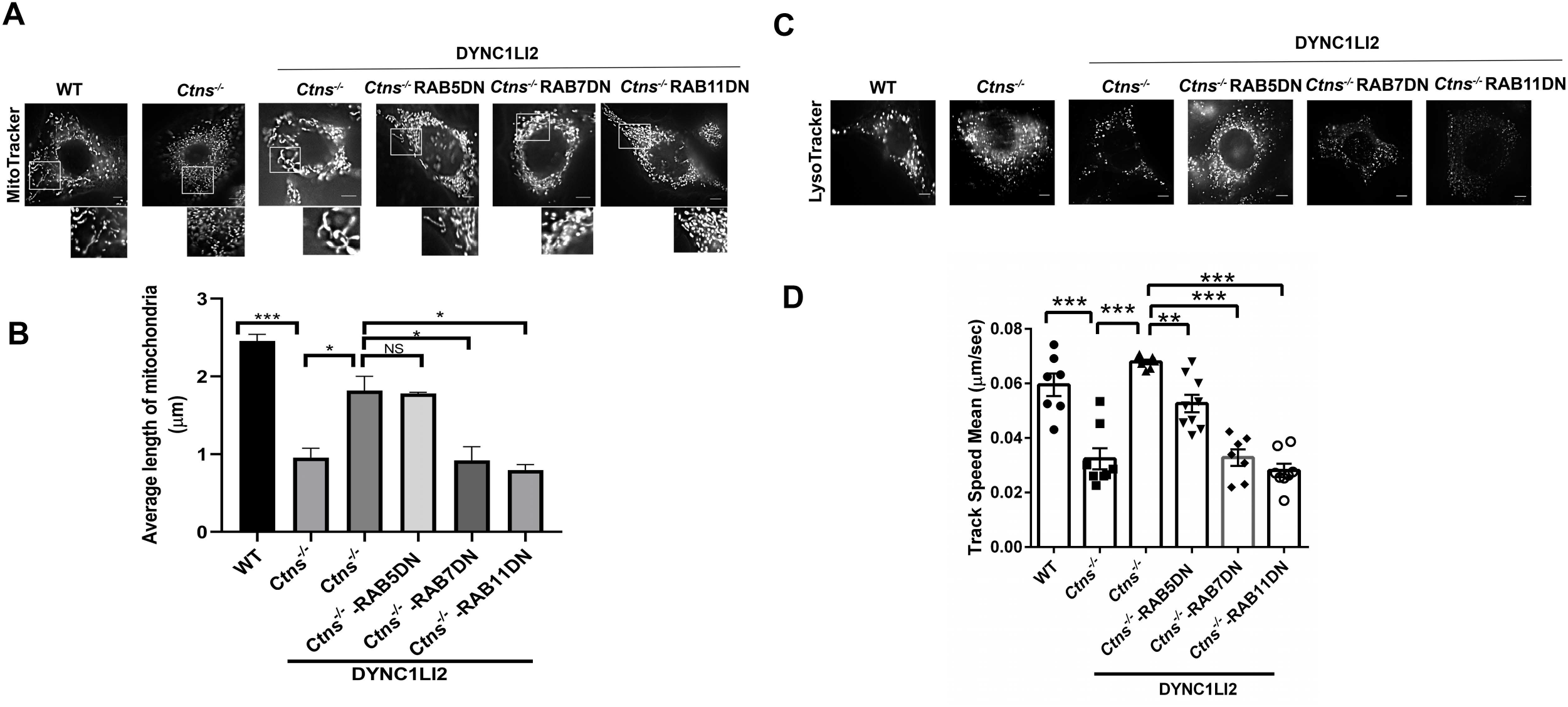
DYNC1LI2-mediated rescue of mitochondrial fragmentation and vesicular trafficking requires RAB7 and RAB11. *A*, TIRF microscopy analysis of WT, *Ctns^-/-^, Ctns*^-/-^ DYNC1LI2 and RAB5DN/RAB7DN/RAB11DN expressing *Ctns*^-/-^ DYNC1LI2 fibroblasts labeled with MitoTracker. *Scale bar*, 6 μm. *B*, Quantification of mitochondrial length. *C*, Representative images of WT, *Ctns^-/-,^ Ctns*^-/-^ DYNC1LI2 and DsRed-RAB5DN/RAB7DN/RAB11DN expressing *Ctns*^-/-^- DYNC1LI2 fibroblasts labeled with LysoTracker. *Scale bar*, 6 μm. *D*, Analysis of vesicle movement. Results are represented as mean ± SEM from at least 7-9 cells per group. The statistically significant differences between the groups are indicated in the figure. *, *p*< 0.05, **, *p*< 0.01, ***, *p*< 0.001. NS, not significant.

### DYNC1LI2-mediated rescue of cystinotic cell function requires the CMA receptor LAMP2A

Trafficking and lysosomal localization of the CMA receptor LAMP2A is mediated by Rab7, its effector RILP, and Rab11^11^. CMA activators increase vesicular trafficking and mediate Rab11 upregulation but LAMP2A knockdown prevents the improvement of cell survival mediated by CMA activators^39^. Here, we analyzed whether DYNC1LI2-dependent rescue of the cystinotic defective phenotype was dependent on LAMP2A. To this end, we analyzed ER stress in cystinotic cells reconstituted with DYNC1LI2, in which LAMP2A expression was downregulated using specific, and previously validated^11^, lentiviral LAMP2A-shRNAs. In **Figures 8 A and B**, we show that the rescue mediated by DYNC1LI2 reconstitution is significantly impaired in cells with decreased expression levels of the CMA receptor LAMP2A. In an independent approach, we analyzed the effect of LAMP2A-downregulation on KDEL expression in cystinotic cells reconstituted with DYNC1LI2 using immunofluorescence analysis of endogenous ER chaperones (**Figs. 8 C and D**). Again, LAMP2A downregulation blocked the ability of DYNC1LI2 to rescue the defective phenotype in cystinosis, thus supporting that LAMP2A and CMA function are necessary for DYNC1LI2 to correct cell function in cystinosis.

**Figure 8.**
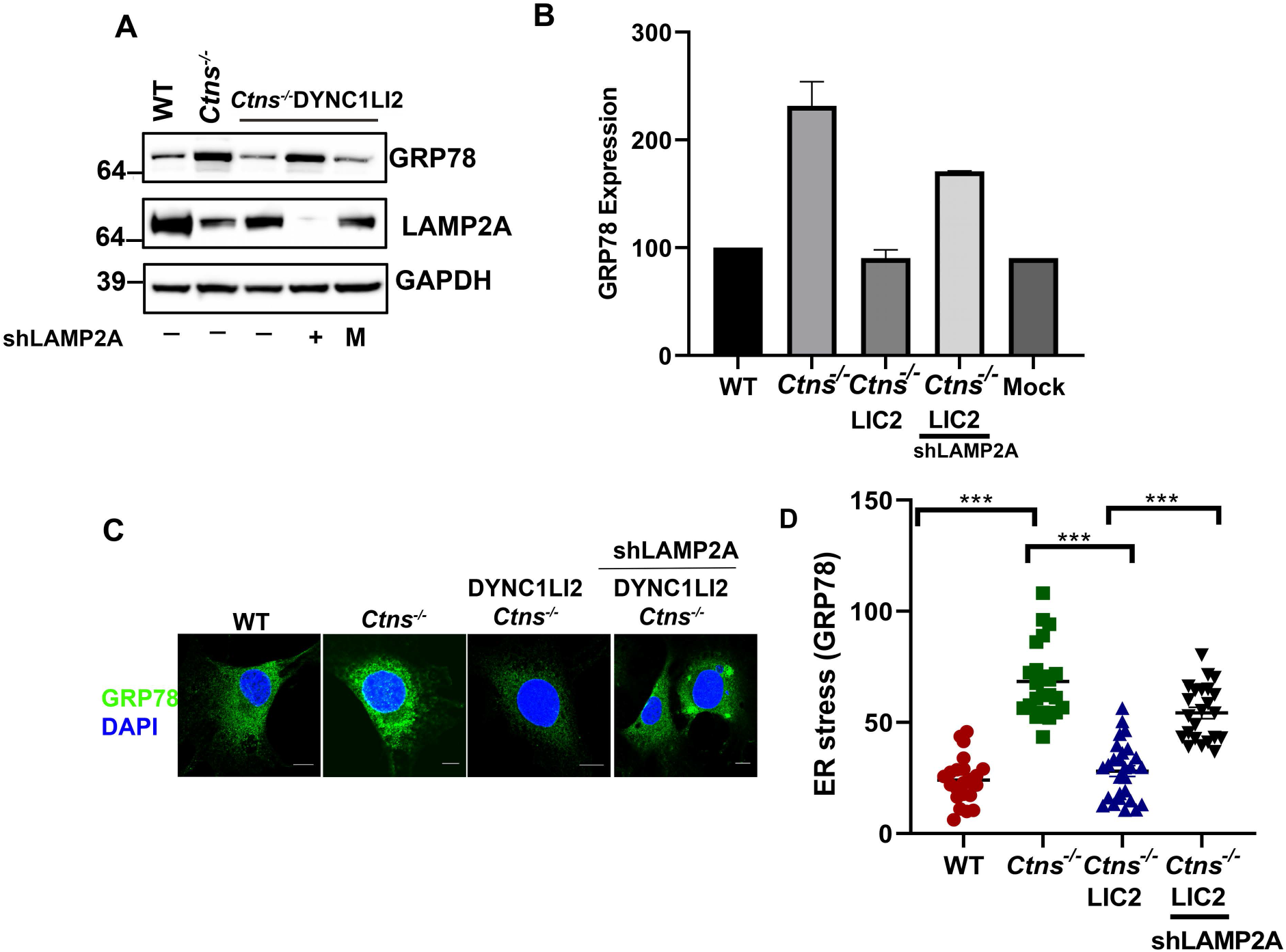
DYNC1LI2-mediated rescue requires the CMA receptor LAMP2A. *A*, Western blot analysis of GRP78 in WT, *Ctns*^-/-^, *Ctns*^-/-^-DYNC1LI2 and LAMP2A-knockdown (shLAMP2A)-expressing *Ctns*^-/-^-DYNC1LI2 fibroblasts. M: mock infected (*Ctns*^-/-^-DYNC1LI2-expressing stable cells infected with empty PLV-Lenti-construct). *B*, Mean ± SEM of GRP78 expression in WT, *Ctns^-/-^, Ctns*^-/-^-DYNC1LI2 and LAMP2A-knockdown (shLAMP2A)-expressing cells. *C*, Immunofluorescence analysis of endogenous ER stress marker expression in wild-type, *Ctns^-/-^, Ctns*^-/-^-DYNC1LI2 (LIC2) and shLAMP2A *Ctns*^-/-^-DYNC1LI2 cells. *Scale bar*, 10 μm. *D*, Quantification of ER stress marker (Grp78) intensity. Mean ± SEM, *\*\*\*p*< 0.001. n=3.

## DISCUSSION

Impaired vesicular trafficking leads to defects in the maintenance of cellular homeostasis but the molecular mechanisms mediating these processes are not well-understood. The Hsc70-dependent translocation of proteins destined for degradation in the lysosome by a process referred to as chaperone-mediates autophagy (CMA), requires the presence of LAMP2A, the only known receptor for CMA, at the lysosomal membrane^34^. Efficient translocation of LAMP2A to the lysosome requires the small GTPases Rab7 and Rab11^40^. The dysregulation of this mechanism is particularly manifested in the lysosomal storage disease cystinosis, in which LAMP2A is retained at Rab11-poitive carrier vesicles with the deleterious effect that the consequent impairment in CMA decreases cell survival^3^. Here, we show that the vesicular trafficking regulator, DYNC1LI2, is downregulated in cystinosis, and its reconstitution corrects the impaired cellular homeostasis by decreasing ER stress and mitochondrial fragmentation. Thus, DYNC1LI2 emerges as a molecular target whose upregulation may improve cellular function in this lysosomal storage disorder.

Because only a subpopulation of lysosomes is active for CMA, LAMP2A lysosomal localization and stabilization at the lysosomal membrane are essential mechanisms to maintain cellular homeostasis. Furthermore, activation of CMA induces the re-localization of CMA-competent lysosomes from the cell periphery to the perinuclear area^46^. In this context, the finding that DYNC1LI2, a protein involved in retrograde transport, regulates LAMP2A lysosomal localization and CMA mediated functions, has important biological significance. Our data showing that DYNC1LI2-mediated rescue of the defective phenotypes observed in cystinosis is attenuated in LAMP2A-downregulated cells establishes a direct link between vesicular trafficking, CMA and cellular homeostasis.

Several Rab GTPases and their effectors have been described to act as molecular links between the dynein molecular machinery and the cargoes they transport. For example, the sorting endosome GTPase, Rab4A, has been implicated in the localization of endosomes to microtubules through the direct binding to DYNC1LI1 in a GTP-dependent manner^24^. The small GTPase Rab11 and its effector Rab11-FIP3 were later shown to regulate the endosomal-recycling compartment through binding to both intermediate-light chains, DYNC1LI1 and DYNC1LI2^32,47^. The Rab7 effector RILP was originally associated with dynactin p150^Glued^, a mechanism that regulates the recruitment of late endosomes to the dynein motor^48^. Later, RILP was shown to recruit late endosomes to dynein motor-associated microtubules through DYNC1LI1 in a dynactin-independent manner^30^. Although downregulation of either DYNC1LI1 or DYNC1LI2 was shown to affect late endosome distribution, based on the more clear association of DYNC1LI1 to late endosomes, it was proposed that DYNC1LI1 plays a more critical role in the endocytic pathway^30^.

Here, we show that inactivation of either Rab11 or Rab7 was sufficient to prevent the rescue of cellular homeostatic mechanisms mediated by DYNC1LI2, thus functionally linking DYNC1LI2 to these GTPases. In principle, this suggest that DYNC1LI2 operates through Rab11 and Rab7 by two essential and independent trafficking mechanisms. While interference with either Rab7 or Rab11 affects the trafficking kinetic parameters of LAMP2A, only Rab7 interference affects the movement of a fast-moving subcellular pool of LAMP2A-positive vesicles^11^. Together, these data suggest that LAMP2A may require the regulation of two sequential mechanisms mediated by Rab11 and Rab7, and that DYNC1LI2 may facilitate this transition. The observation that DYNC1LI2 preferentially upregulates the dynamics of highly motile endo-lysosomes, Rab7-expressing vesicles, further support this hypothesis. Alternatively, Rab7 and Rab11 may define two independent sub-types of LAMP2A-positive vesicles but both pools should be necessary to translocate competent amounts of LAMP2A to lysosomal membranes.

Both DYNC1LI1 and DYNC1LI2 bind to the dynein motor heavy chain as well as to Rab11 and Rab7 effectors. However, only DYNC1LI2 but not DYNC1LI1 is downregulated in cystinosis and expression of DYNC1LI2 but not DYNC1LI1 rescues the defective phenotypes observed in cystinotic cells. This suggests that specific molecular determinants in addition to motor subunits and Rab adaptors are necessary for DYNC1LI2 to specifically exert its biological function in cystinotic cells. Of note, the interaction of DYNC1LI1 but not DYNC1LI2 with Rab effectors is regulated by post-translational modification including phosphorylation^45^ and so, it is possible that this type of regulation is absent in cells where DYNC1LI2 is predominantly active.

In conclusion, we show that DYNC1LI2 but not DYNC1LI1 plays a fundamental role in the translocation of LAMP2A to CMA-competent lysosomes and identified LAMP2A and CMA as a novel mechanism modulated by DYNC1LI2. This mechanism is directly associated with the maintenance of important cellular homeostatic processes in a Rab7- and Rab11-dependent manner. Thus, DYNC1LI2 upregulation leads to the correction of defective phenotypes including endoplasmic reticulum stress, increased mitochondria fragmentation and decreased cell survival. These conditions are particularly manifested in cystinotic cells, thus highlighting a role for DYNC1LI2 in vesicular trafficking in this disease. Because upregulation of DYNC1LI2 leads to the correction of cellular homeostatic mechanisms in proximal tubule cells, DYNC1LI2 emerges as a new possible target to correct pathological processes specific for kidney disfunction in this lysosomal storage disorder.

## Materials and methods

### Animal models

All animal studies were performed in compliance with the United States Department of Health and Human Services Guide for the Care and Use of Laboratory Animals. All studies were conducted according to National Institutes of Health and institutional guidelines and with approval of the animal review boards at The Scripps Research Institute and University of California, San Diego. The C57BL/6 *Ctns*^-/-^ mice were described before^49^.

Neonatal mouse embryonic fibroblasts from *Ctns*^-/-^ and wild-type mice were prepared by standard procedures^3^. The cystinosin knockout human proximal tubule cell line was generated using CRISPR/Cas9 and described previously^39^. Neonatal murine fibroblasts, proximal tubule cell line and 293FT cells (ATCC) were maintained in Dulbecco’s modified Eagle’s medium (Gibco) supplemented with 10% FBS (Corning Cellgro) and penicillin/streptomycin/glutamine (Life Sciences).

### Gene expression Microarray analysis

Animals were housed and studied according to NIH Guidelines for the Care and Use of Laboratory Animals. Sixteen months old C57BL/6 wild-type (n=6) and *Ctns*^-/-^ (n=6) mice were euthanized and the kidneys immediately removed and stored at −80°C in RNA Later (Life Technologies). Tissues were subsequently ground using Precellys 24 (Bertin Technologies) and RNA was isolated using the Qiagen AllPrep DNA/RNA Mini kit (Qiagen). The RNA was run on a Bioanalyzer (Agilent) for quantification and quality assessment. The Ambion WT Expression Kit was used to generate double-stranded biotinylated cDNA and the Affymetrix HT WT Terminal Labeling Kit was used to prepare the cDNA for hybridization to Affymetrix GeneChip Mouse Gene 1.1 ST arrays (Affymetrix). The double-stranded biotinylated cDNA from each tissue for each mouse was run on individual Affymetrix GeneChip. The data was collected as CEL files and quality control analysis performed with Affymetrix Expression Console. Normalized signal intensities were generated with Robust Multichip Average (RMA) which employs a background adjustment and quantile normalization strategy^50^. Genes without an average log2 transformed signal above 6 in either the wild-type or *Ctns*^-/-^ group were removed from further analysis. Class comparisons of variance by two-way t-tests for two sample comparisons (p<0.001) were performed using BRB-ArrayTools (http://linus.nci.nih.gov/BRB-ArrayTools.html) to identify the set of significantly differentially expressed genes between wild-type and *Ctns*^-/-^ mice in each tissue.

### Constructs, transfections, and transductions

Lentiviral vectors expressing pLVXGFP-DYNC1LI2 (plasmid #66601), pLVXGFP-DYNC1LI1 (plasmid # 66600), were obtained from Addgene (deposited by Dr. David Stephens) and were analyzed by sequencing for confirmation (Retrogene). The pLVGFP expression vector was used as a control in mock infections. 293FT cells were transfected with the pLV constructs along with packaging plasmids pVSVG (Clontech) and pLV-CMV-delta 8.2 (Massey, A.C 2006) using the Lipofectamine LTX transfection reagent (Life Technologies). Virus-containing supernatants were collected at 24 and 48 h post transfection, filtered, and then used to infect cells without additional processing or concentration. Wild-type and *Ctns*^-/-^ type cells of murine fibroblasts and PTC cells were transduced with lentiviruses in the presence of 5 μg/ml Polybrene. After 24 h, the virus-containing medium was replaced with normal growth medium. The transduction efficiency obtained using this procedure was > 90%^12^. To make stable cell lines, infected cells were kept under selection pressure for puromycin, selected by GFP-expression and analyzed by WB to select cells that maintained stable levels of expression. The constructs DsRed-Rab5DN, DsRed-Rab7DN were described previously^11^. DSRed-Rab11DN was purchased from Addgene^11,51^. DYNC1LI2 expressing *Ctns*^-/-^ MEF cells and 293FT cells were transfected using Lipofectamine 3000 (Thermo Fisher Scientific, 2103388) following the manufacturer’s instructions. Lentiviral mouse LAMP2A shRNA was transduced in DYNC1LI2-*Ctns*^-/-^ cells also as described before^11^.

### Gel electrophoresis and immunoblotting

Cells were lysed in lysis buffer (20 mM Tris (pH 7.4), 150 mM NaCl, and 1% Triton X-100) in the presence of protease/inhibitor (Roche Applied Science) and phosphatase inhibitors (Calbiochem). Protein concentration was measured using the colorimetric Bradford assay (Bio-Rad, 5000006). Gel electrophoresis was carried out using 4–12% gradient gels or 12% Nu-PAGE gels (Life Sciences). Proteins were transferred onto 0.45-μm nitrocellulose membranes and the membranes were incubated overnight at 4°C in the indicated primary antibodies, followed by incubation with HRP-conjugated secondary antibodies. The blots were developed using SuperSignal West Pico or Femto chemiluminescence substrate systems (Thermo Fisher Scientific). Proteins were visualized using Azure Biosystems technology. The following antibodies were used for immunoblotting in this study: rabbit anti-DYNC1LI2 (PA5-45861), anti-DYNC1LI1 (PA5-31644), anti-Rab5 (2143T), anti-Rab7 (9367S), anti-Rab11a (5589S), anti-actin (SC7210); anti-GAPDH (GT239); anti-GFP (SC9996), anti-FLAG (Genecopoeia) and anti-KDEL (Enzo Life Sciences, ADI-SPA-827).

### Immunofluorescence, immunohistochemistry, and confocal analysis

Murine fibroblast and PTCs were seeded at 70% confluence in a 4-chamber 35-mm glass-bottom dish (In Vitro Scientific, D35C4-20-1.5-N), then fixed with 4% paraformaldehyde for 8 min and blocked with 1% BSA in PBS, in the presence of 0.01% saponin, for 1 h. Samples were labeled with the indicated primary antibodies overnight at 4°C in the presence of 0.01% saponin (Calbiochem, 558255) and 1% BSA. Samples were washed 3 times and subsequently incubated with the appropriate combinations of Alexa Fluor (488 or 594)-conjugated anti-rabbit, anti-rat, or antimouse secondary antibodies (Thermo Fisher Scientific, A-21206; A-21207; A-21208; A-21209; A-21202; A-21203, respectively). Nuclei were stained with 4,6 - diamidino-2 phenylindole (DAPI) and samples were preserved in Fluoromount-G reagent (SouthernBiotech) and kept at 4°C until analyzed. Samples were analyzed with a Zeiss LSM 710 or Zeiss LSM 880 laser-scanning confocal microscope attached to a Zeiss Observer Z1 microscope at 21°C, using a 63x oil Plan Apo, 1.4-numerical aperture objective. Images were collected using ZEN-LSM software and processed using ImageJ and Adobe Photoshop CS4. The exposure time and gain were maintained throughout the experiment to comparatively analyze wild-type and *Ctns*^-/-^ cells. Analysis and quantification of protein colocalization were performed using ZEN-LSM software. The following antibodies were used for immunofluorescence in this study: anti-LAMP1 (Santa Cruz Biotechnology, sc-19992); anti-LAMP2A (Abcam, ab18528 or ThermoFischer Scientific 51-2200), rabbit anti-GFP (3085S) and anti-KDEL (Enzo Life Sciences, ADI-SPA-827).

### TIRF microscopy

Cells labelled with LysoTracker and MitoTracker were transferred to a microscopic stage and maintained at 37°C. Pseudo-TIRF microscopy (oblique illumination) experiments were performed using a X100 1.45 numerical aperture TIRF objective (Nikon) on a Nikon TE2000U microscope custom-modified with a TIRF illumination module as described^11^. Images were acquired on a 14-bit cooled charge-coupled device camera (Hamamatsu) controlled through NIS-Elements software. For live cell imaging, the images were recorded using 200–500-ms exposures depending on the fluorescence intensity of the sample. The track speed mean was then analyzed using Imaris as described before^11,39^. Wild-type and *Ctns*^-/-^ cells were analyzed comparatively by maintaining exposure time and gain throughout the experiment. Length of mitochondria were analyzed using the ‘Measure’ tool in ImageJ software.

### MTT Assay

MTT assays were performed as a measurement of cell metabolism and survival to oxidative stress insult. MEFs were seeded at 4,000 cells/well onto 96-well plates. After 48 h, the cells were replenished with fresh culture medium containing increasing concentrations (0-500 μm) of H_2_0_2_ for 16 h. After treatment, cell viability was determined by a modified version of the MTT assay. The assay was carried out by discarding the cell culture medium and replace with 100 μl of fresh culture medium containing 5.0 mg/ml MTT for 4 h at 37°C. Next, the cells were solubilized with 100 μl of a solution containing 50% dimethylformamide and 20% SDS (pH: 4.7), for 16 h. Absorbance (560) nm was measured using a microplate reader (SpectroMax Gemini; Molecular Devices Corp., Sunnyvale,CA).

### Statistical analysis

Data are presented as mean, and error bars correspond to standard errors of the means (SEMs) unless otherwise indicated. Statistical significance was determined using the unpaired Student’s t-test using GraphPad InStat (version 3) or Excel software, and graphs were made using GraphPad Prism (version 6) software.

## Acknowledgements

Supported by fellowships from the Cystinosis Research Foundation to FR and JZ, by the National Institutes of Health grant numbers: R01HL088256, R01AR070837, R01DK110162 and R21EY028642 to S.D.C, and R01-DK090058 and R01-NS108965 for S.C.

## Notes

### Competing Interest Statement

The authors have declared no competing interest.

